# Karyotype asymmetry in *Cuscuta* L. subgenus *Pachystigma* reflects its repeat DNA composition

**DOI:** 10.1101/2021.08.09.455742

**Authors:** Amalia Ibiapino, Mariana Báez, Miguel A. García, Mihai Costea, Saša Stefanović, Andrea Pedrosa-Harand

## Abstract

*Cuscuta* is a cytogenetically diverse genus, with karyotypes varying 18-fold in chromosome number and 89-fold in genome size. Each of its four subgenera also presents particular chromosomal features, such as bimodal karyotypes in *Pachystigma*. We used low coverage sequencing of the *Cuscuta nitida* genome (subgenus *Pachystigma*), as well as chromosome banding and molecular cytogenetics of three subgenus representatives, to understand the origin of bimodal karyotypes. All three species, *C. nitida, C. africana* (2*n* = 28) and *C. angulata* (2*n* = 30), showed heterochromatic bands mainly in the largest chromosome pairs. Eighteen satellite DNAs were identified in *C. nitida* genome, two showing similarity to mobile elements. The most abundant were present at the largest pairs, as well as the highly abundant ribosomal DNAs. The most abundant Ty1/Copia and Ty3/Gypsy elements were also highly enriched in the largest pairs, except for the Ty3/Gypsy CRM, which also labelled the pericentromeric regions of the smallest chromosomes. This accumulation of repetitive DNA in the larger pairs indicates that these sequences are largely responsible for the formation of bimodal karyotypes in the subgenus *Pachystigma*. The repetitive DNA fraction is directly linked to karyotype evolution in *Cuscuta*.

**Highlights:** *Cuscuta* subgenus *Pachystigma* contains species with strikingly bimodal karyotypes. The emergence of these karyotypes is linked to the enrichment of varied repetitive sequences in the largest chromosomal pairs.

## Introduction

Most angiosperms have symmetric karyotypes (Stebbins, 1971; Weiss-Schneeweiss and Schneeweiss, 2013), with chromosomes similar in size and morphology. However, several lineages have been known to have asymmetrical karyotypes, characterized by centromeres at different positions along chromosomes, or chromosomes of different sizes. Karyotypes with two sets of chromosomes markedly different in size are called bimodal and represent the extreme of karyotype asymmetry (Stebbins, 1971; McKain *et al*., 2012). Three main hypotheses were suggested for the origin of bimodal karyotypes. The first mechanism involves chromosomal rearrangements such as fusion-fission events. Thus, larger chromosomes can be the product of small chromosome fusions or small chromosomes the result of fission of larger ones (Schubert and Lysak, 2011). *Lygosoma bowringii* Günther 1864, a lizard of the Scincidae family, has 2*n* = 32, with 18 macrochromosomes and 14 microchromosomes. As in most reptiles, macrochromosomes have originated from the fusion of microchromosomes (Lisachov *et al*., 2018, 2020). The second mechanism is allopolyploidy involving parental species with different chromosomal sizes, suggested for some genera such as *Agave* L. (McKain *et al*., 2012). The third possibility is the progressive amplification of repetitive DNA sequences in one set of chromosomes (de la Herrán *et al*., 2001). In *Eleutherine* Herb. (Iridaceae), for example, an accumulation of different families of repetitive sequences in the larger chromosome pair was suggested to be the cause of the differences between both chromosome sets (Báez *et al*., 2019).

Repetitive DNA constitutes a large fraction of plant genomes and can be found either organized in tandem (micro-, mini- and satellite DNAs), or dispersed through the genome (transposons and retrotransposons) (Heslop-Harrison and Schwarzacher, 2011). Transposable elements are capable of moving within the genome, impacting genome structure and even the function of genes (Bourque *et al*., 2018). The highly abundant repetitive sequences are frequently associated with heterochromatin formation located at the (peri-) centromeres, subtelomeres and in interstitial heterochromatic blocks (Barros e Silva *et al*., 2010; Van-Lume *et al*., 2019). The effects of transposable element and satellite DNA accumulation on genomes are dynamic and can lead to significant increase in genome size. In species of the genus *Zea* L., the accumulation of repetitive DNAs families, mainly LTR retrotransposons like Ty3/Gypsy, resulted in a two-times larger genome in *Zea luxurians* (Durieu) R.M.Bird in relation to *Z. mays* L. and *Z. diploperennis* Iltis, Doebley & R. Guzmán in less than two million years (Estep *et al*., 2013).

Satellite DNAs are composed by monomers that are oriented head-to-tail and can vary in length, nucleotide composition, sequence complexity, and abundance. These sequences frequently form clusters that can rapidly change in number, position and size (Garrido-Ramos, 2015; Biscotti *et al*., 2015). A mutation that occurs within a monomer can spread among the repeat units or be eliminated by homogenization (Plohl *et al*., 2012). The mechanism of concerted evolution, for example, can generate varied patterns of repetitive DNA families, producing in general homogeneity within species and diversity between species. Thus, different species can have different families of satellite DNAs or these satellites can be shared between related species (Feliner and Rosselló, 2012; Plohl *et al*., 2012). Furthermore, these tandem repeats can be species-specific or even chromosome specific. Some tandem repeats are highly conserved among species, such as the 5S and 35S ribosomal DNAs (rDNAs), which encode for the ribosomal RNAs. But most repetitive families are usually non-coding sequences evolving rapidly and generating genomic differentiation (Biscotti *et al*., 2015).

The parasitic genus *Cuscuta* L. (Convolvulaceae Juss.) includes some 200 species, divided into four subgenera: *Grammica* (Lour.) Peter, Engl. & Prantl, *Pachystigma* (Engelm.) Baker & C.H. Wright, *Cuscuta* Yunck, and *Monogynella* (Des Moul.) Peter, Engl. & Prantl (García *et al*., 2014; Costea *et al*., 2015a). Subgenus *Grammica*, with about 150 species, has almost exclusive distribution in the Americas. *Pachystigma* includes only five species, all endemic to South Africa, and *Cuscuta* is native to Europe, Africa, and Asia, with a few species introduced and naturalized in the Americas, Australia, and New Zealand. Subgenus *Monogynella* had its origin in Central Asia from where it dispersed to S, E and SE Asia, Europe, Africa, and one species, *C. exaltata* Engelm., is native to south-eastern North America (García *et al*., 2014; Costea *et al*., 2015*b*).

The genus *Cuscuta* shows high cytogenetic variation in chromosome number (2*n* = 8 to 2*n* = 150), chromosome size (1.66 μm to 21.60 μm), and genome size (1C = 0.39 Gbp to 1C = 34.73 Gbp). The genus also presents symmetric to bimodal karyotypes, as well as monocentric and holocentric chromosomes (García and Castroviejo, 2003; Guerra and García, 2004; McNeal *et al*., 2007; Ibiapino *et al*., 2019, 2020; García *et al*., 2019; Oliveira *et al*., 2020; Neumann *et al*., 2020). Each *Cuscuta* subgenus seems to have different karyotypic features. Species of subgenus *Monogynella* have the largest genome sizes and the largest chromosomes. Subgenus *Cuscuta* is the only one that has species with exclusively holocentric chromosomes. Subgenus *Grammica* presents the largest variation in chromosome number and size. This subgenus has at least five cases of interspecific hybridization which can contribute to this chromosome number variation (Fogelberg, 1938; Pazy and Plitmann, 1994; García, 2001; García and Castroviejo, 2003; McNeal *et al*., 2007; Ibiapino *et al*., 2019; García *et al*., 2019). A preliminary study of two species of *Pachystigma* revealed bimodal karyotypes and extensive heterochromatic blocks in the larger chromosomes, suggesting the influence of repetitive DNA in the emergence of bimodality in this subgenus (García *et al*., 2019). An asymmetrical karyotype was also reported for some populations of the holocentric *C. epithymum* (L.) L. (subgenus *Cuscuta*), with 2*n* = 14 individuals showing bimodal karyotype while 2*n* = 16 individuals having symmetric karyotypes (García and Castroviejo, 2003).

Species of the genus *Cuscuta* also vary in heterochromatin content, ranging from species with few bands and few rDNA sites, such as *C. denticulata* (Ibiapino *et al*., 2019), up to species with numerous bands, where heterochromatin may have contributed to the expansion of the genome size, as in *C. monogyna* and *C. indecora* (Ibiapino *et al*., 2020; Oliveira *et al*., 2020). In the latter two species, heterochromatin may have contributed to maintaining karyotype symmetry, since both have similar karyotypes, but belong to different subgenera (Ibiapino *et al*., 2020). Repeat DNA composition was investigated in 12 *Cuscuta* species, demonstrating that the extensive variation in genome size in species of this genus is caused by the differential accumulation of repetitive sequences (Neumann *et al*., 2020). However, no representatives of subgenus *Pachystigma* were included in that study.

Our current work investigates heterochromatin distribution in three of the five species of the subgenus *Pachystigma* (*C. nitida* E. Mey. ex Choisy, *C. africana* Thunb. and *C. angulata* Engelm.) and evaluates the repetitive DNA composition of *C. nitida* genome, in order to better understand the role played by repetitive DNA sequences in the emergence of bimodal karyotypes within this subgenus.

## Materials and methods

### Material

Flower buds of two accessions of *C. africana*, one of *C. angulata* and three of *C. nitida* (subgenus *Pachystigma*) were collected in November 2017 from the Cape region of South Africa, where they are endemic (Table 1). Vouchers were deposited at the herbaria of the University of Toronto Mississauga (TRTE) and Wilfrid Laurier University (WLU), Canada.

**Table 1:**
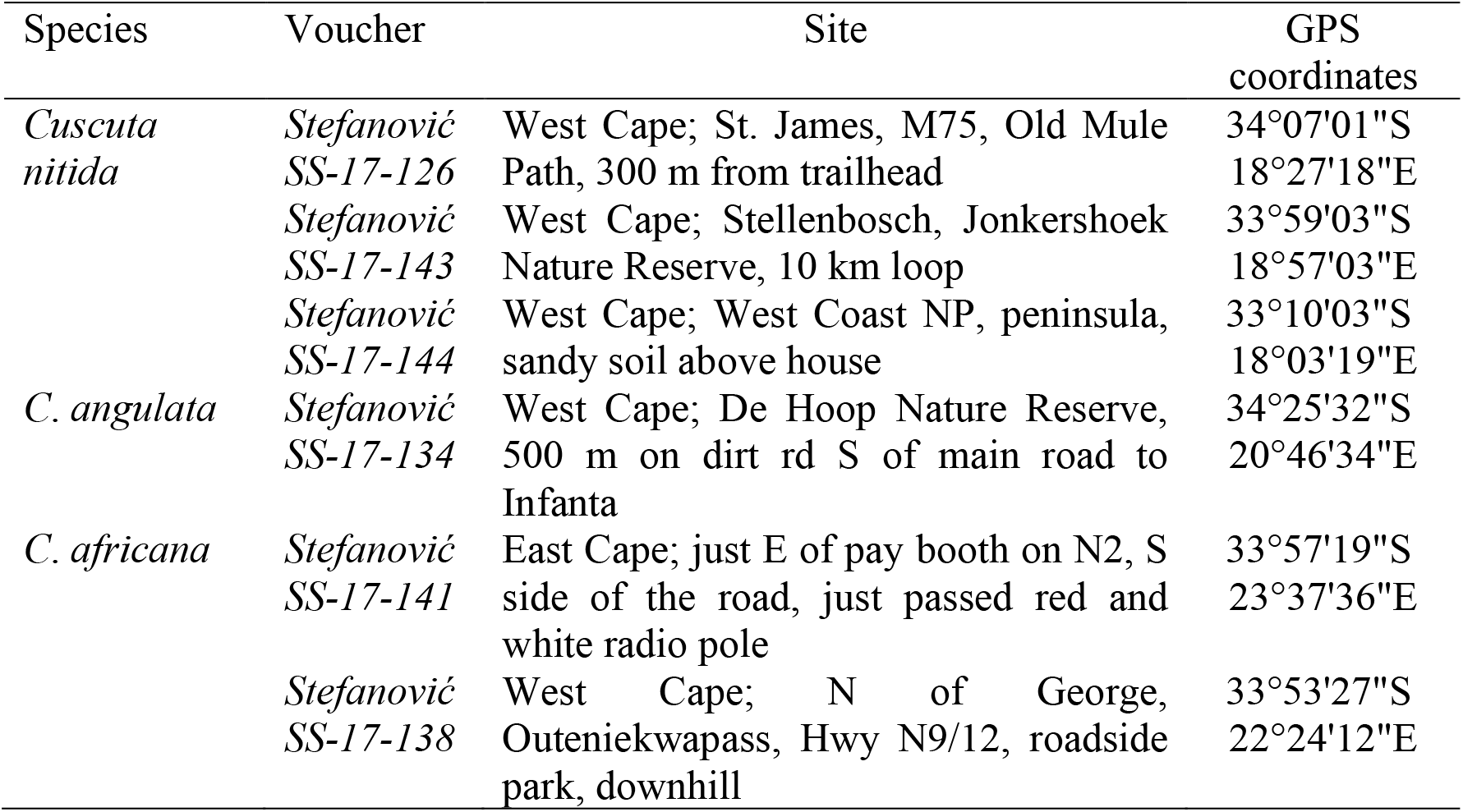
Collection sites for three *Cuscuta* species of the subgenus *Pachystigma*

### Slide preparation and CMA/DAPI double staining

Slides were prepared using flower buds collected and fixed in the field in Carnoy (ethanol: acetic acid, 3:1, v/v). The material was washed in distilled water, digested in an enzymatic solution containing 2% cellulase (Onozuka) and 20% pectinase (Sigma) for 40 minutes. The slides were prepared by air drying, mainly using the ovary wall, as described by De Carvalho and Saraiva (1993), with small modifications. After the material was macerated and dried, the slides were dipped in 60% acetic acid for up to 5 minutes to clear the cytoplasm. Finally, the slides were left at 37° C until completely dry.

For double CMA/DAPI staining, the slides were aged at room temperature for three days, stained with 8 μL of 0.1 mg/μL chromomycin A3 (CMA) for 60 minutes, mounted in 8 uL of 1μg/mL 4’, 6-diamidino-2-phenylindole (DAPI) in mounting medium (glycerol:McIlvaine buffer pH 7.0, 1:1, v/v), and aged again for three days at room temperature. The images were captured with a COHU CCD camera attached to a Leica DMLB fluorescence microscope equipped with Leica QFISH software. After image capture, slides were destained for 30 minutes in Carnoy, for one hour in absolute ethanol and stored at −20° C for *in situ* hybridization.

For chromosomal measurements, five metaphases of *C. nitida* were used. Chromosomes were measured with the ruler tool in Adobe Photoshop CS3 version 10.0.

### DNA extraction and *in silico* repetitive DNA analysis

*Cuscuta nitida* genomic DNA was extracted following Doyle and Doyle (1987) protocol. Sequencing of the total genomic DNA generated low coverage (0.01×), 250-bp paired-end reads in an Illumina HiSeq 2500 (BGI, Hong Kong, China). Repetitive DNA analysis was performed by the RepeatExplorer pipeline (https://galaxy-elixir.cerit-sc.cz/; Novak *et al*., 2013), where reads showing at least 95% similarity in at least 55% of its length were clustered together.

Clusters showing an abundance greater than 0.01% were automatically annotated and manually checked. Clusters similar to plastomes or mitogenomes were considered putative contamination and excluded from the final annotation. All contigs with tandem repetitions identified by TAREAN (Novák *et al*., 2017), as well as other satellites not identified by this tool, but which presented typical satellite graph layouts after clustering, were confirmed with DOTTER (Sonnhammer and Durbin, 1995). High abundance dispersed elements had their integrase domain identified using the NCBI Conserved Domain Search (https://www.ncbi.nlm.nih.gov/Structure/cdd/wrpsb.cgi). The consensus sequences of the satellites and the integrase domains of the transposable elements were used for primer design using the primer design tool implemented in Geneious version 7.1.9 (Kearse *et al*., 2012).

The consensus sequences of all identified satellites were compared in order to verify their homology. The consensus monomers that showed similarity in DOTTER were aligned using Muscle in Geneious. Different satellite families were considered as part of the same superfamily when monomer sequences showed identity between 50% and 80%. Sequences with 80-95% similarity were considered subfamilies of the same family and similarity greater than 95% were considered variants of the same family (Ruiz-Ruano *et al*., 2016) As two of these satellites showed similarity with transposable elements, alignments were made of the consensus satellite sequence with the most similar transposable element domains indicated by the RepeatExplorer. One of the satellites that showed similarity with transposable elements also showed *in situ* colocalization with the 35S rDNA cluster. Therefore, a comparison of the satellite consensus sequence with a putative *C. campestris* (GenBank accession number PRJEB19879) 35S rDNA consensus sequence, assembled using the NOVOPlasty algorithm (Dierckxsens *et al*., 2017) was included. This assembly was made using Illumina reads obtained from Vogel *et al*., (2018). After assembled, the complete 35S rDNA was aligned with the satellite consensus sequence using Muscle in Geneious. Satellites were named as follows: code referring to the species name (Cn), followed by “Sat”, a number referring to the abundance order, and the size of the consensus monomer in base pairs.

### Repeat amplification, probe preparation and *in situ* hybridization (FISH)

Polymerase chain reaction (PCR) for repeat amplification was performed in 50 μL reactions containing 200 ng of *C. nitida* genomic DNA, 1× PCR buffer (20 mM Tris-HCl pH 8.4, 50 mM KCl), 2 mM MgCl_2_, 0.1 mM dNTPs, 0.4 μM of each primer, 0.4× TBT (750 mM trehalose, 1 mg/ml BSA, 1% Tween 20, 8.5 mM Tris hydrochloride) and 0.6 μL of a homemade Taq Polymerase. Amplification program was 1× 94°C for three minutes, plus 30 cycles of 94 °C for one minute, 55-65 °C for one minute (see Supplementary Table 1 for annealing temperatures of each primer pair) and 72 °C for one minute, followed by a final extension of 7 min at 72 °C. Primers for amplification of satellite superfamilies SF1 and SF2 annealed in a conserved region shared between the satellite variants from each superfamily.

The PCR products were sequenced to confirm its identity and labelled in a total volume reaction of 12.5 μL, containing 1 μg of amplified DNA, 1× Nick Translation buffer (0.5 M Tris HCl pH 7.5; 50 mM MgCl_2_), dNTP mix (0.016 mM each of dATP, dCTP, dGTP), 0.08 mM Cy3-dUTP or Alexa-dUTP, 7,5 U of DNA Polymerase I and 0,006 U of DNase I. The mixture was incubated at 15° C for one hour or longer if needed, until most fragments were under 500 bp, and reactions were stopped using 0.5 M EDTA.

For rDNA probes, the plasmids D2 of *Lotus japonicus* (Regel) K. Larsen (5S rDNA) and *pTa71* of wheat (25-28S, 5.8S and 18S rDNA) were used (Pedrosa *et al*., 2002; Sousa *et al*., 2011). Probes were labelled by Nick translation with Cy3-dUTP (5S), as described above, and digoxigenin 11-dUTP (35S) with a Nick Translation kit (Invitrogen - Oregon, USA).

Fluorescence *in situ* hybridizations followed Pedrosa *et al*. (2002). The hybridization mixture, composed of 50% formamide, 10% dextran sulphate, 2× SSC, and 5 ng/μl probe, was denatured at 75°C for 10 minutes. Slides were denatured for 5 minutes with the hybridization mixture and hybridized for 18-20 hours at 37°C in a humid chamber. Final stringency was 76% for 5S and 35S rDNA and satellite DNAs. The 35S rDNA probe was detected with anti-digoxigenin produced in sheep, conjugated with FITC (Roche - Basel, Switzerland) and the signal amplified with anti-sheep IgG produced in rabbit conjugated with FITC (Serotec - California, EUA). The slides were mounted as described above. The transposable elements were hybridized with low stringency (40%), similar to Ribeiro *et al*. (2017a).

To verify the putative localization of telomeric sequences at interstitial chromosome sites, the ND-FISH protocol described by Cuadrado *et al*., (2009) was applied. Thirty μL of the hybridization solution containing 2 pmol (25 ng) of the diluted probe (TTTAGGGTTTAGGGTTTAGGGTTTAGGGT5 directily labelled with Cy3, Macrogen, Seoul, Korea) in 2× SSC was added per slide and cover with a coverslip. The slide was incubated 2 h at room temperature protected from light. The coverslip was removed with 2× SSC, washed in 4× SSC/0.2% Tween 20 at room temperature for 10 minutes under agitation, and mounted in DAPI with mounting medium as described above. The satellites and rDNA images were captured as previously described. For transposable elements, images were captured using an epifluorescence Leica DMLB microscope equipped with a COHU 4912-5010 CCD Camera using the Leica QFISH software.

## Results

### Bimodal karyotypes are a typical feature of the subgenus *Pachystigma*, with large chromosomes enriched in heterochromatin

All three samples of *C. nitida* analysed showed 2*n* = 28 and bimodal karyotypes (4L+24S), with two larger chromosomes pairs (average sizes of 12.34 μm and 8.19 μm) and 12 smaller chromosome pairs (average size of 2.67 μm). The second largest pair harbours a proximal nucleolus organizing region (NOR), as evidenced by the large decondensed region between both chromosome arms. The total haploid complement size was 107.6 μm.

For heterochromatin characterization, CMA/DAPI double staining was performed in the three species. *Cuscuta nitida* showed only two pairs of heterochromatic bands, a large CMA^+^/DAPI^-^ band in the short arm of the largest chromosome pair (Fig. 2 A-C) and a second CMA^+^/DAPI^-^ band colocalised with the NOR in the second chromosome pair (Fig. 2 B-C). *In situ* hybridization with 5S and 35S rDNA revealed one large pair of 5S site, co-localized with the CMA^+^ band at the short arm of the largest pair (Fig. 3 A-B). One major and one minor pair of 35S rDNA sites were observed, both in the large chromosome pairs. The major 35S site was observed highly decondensed, co-localised with a CMA^+^ band in the second largest chromosome pair, while the minor site, not always visible, was observed proximally at the largest pair (Fig. 3 B-D).

*Cuscuta africana* and *C. angulata* also showed bimodal karyotypes (Fig. 2 D-I). *Cuscuta africana* exhibited karyotype similarities with *C. nitida* (2*n* = 28, 4L +24S), but the heterochromatic band present in the largest pair was DAPI^+^/CMA^-^ (Fig. 2 D-F). On the other hand, *C. angulata* (2*n* = 30, 10L+20S) presented ten large chromosomes in its karyotype. These chromosomes have numerous heterochromatic bands, mainly DAPI^+^/CMA^-^. In addition, the set of smaller chromosomes of this species has pericentromeric bands, mainly DAPI^+^/CMA^-^ (Fig. 2 I). Thus, the heterochromatin characterization evidenced the presence of bimodal karyotypes in the three analysed species of *Pachystigma*, with a high number of heterochromatic bands in the large chromosome pairs, with different compositions, GC or AT rich, depending on the species.

### The repetitive fraction of *C. nitida* genome is rich in tandem repeats

To understand the composition of the repetitive DNA fraction and its relation to the heterochromatin content, we performed genome skimming in *C. nitida* and characterized the most abundant DNA repeats. A total of 5,156,846 reads were generated, of which 1,173,600 reads were randomly sampled by RepeatExplorer for analysis. A total of 53,117 clusters were identified and 330 clusters, containing at least 0.01% of genome abundance, were grouped into 322 superclusters (Fig. S1) and annotated. Five and 33 clusters that showed similarity to mitochondrial and plastid sequences, respectively, were excluded from further analysis (Table 2).

**Table 2:**
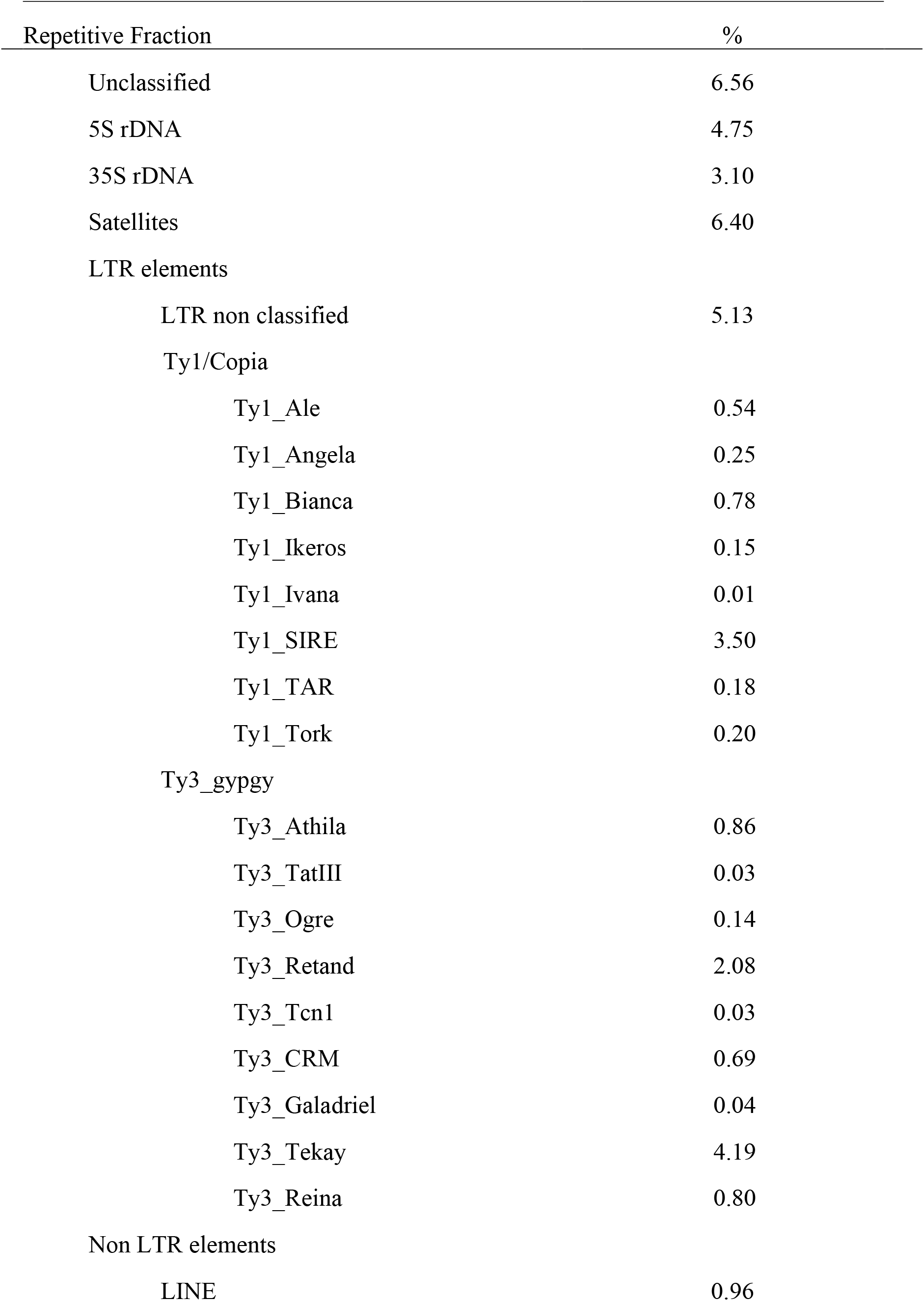
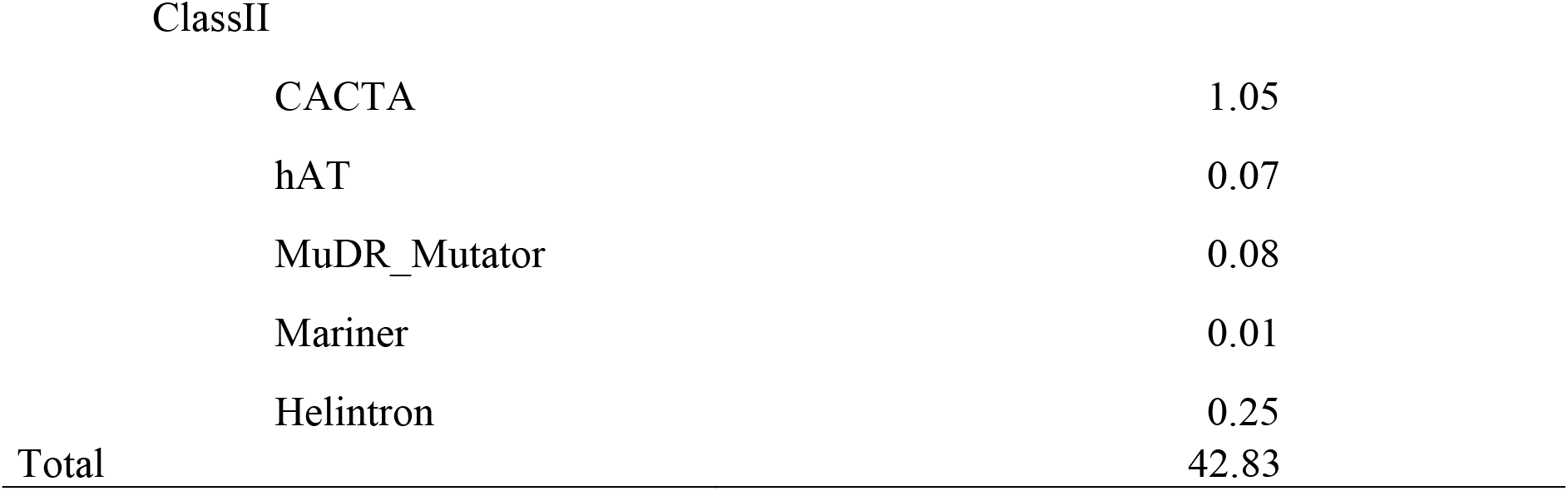
General annotation of the repetitive fraction of *Cuscuta nitida*

| Repetitive Fraction | % |
| --- | --- |
| Unclassified | 6.56 |
| 5S rDNA | 4.75 |
| 35S rDNA | 3.10 |
| Satellites | 6.40 |
| LTR elements |  |
| LTR non classified | 5.13 |
| Ty1/Copia |  |
| Ty1_Ale | 0.54 |
| Ty1_Angela | 0.25 |
| Ty1_Bianca | 0.78 |
| Ty1_Ikeros | 0.15 |
| Ty1_Ivana | 0.01 |
| Ty1_SIRE | 3.50 |
| Ty1_TAR | 0.18 |
| Ty1_Tork | 0.20 |
| Ty3_gypgy |  |
| Ty3_Athila | 0.86 |
| Ty3_TatIII | 0.03 |
| Ty3_Ogre | 0.14 |
| Ty3_Retand | 2.08 |
| Ty3_Tcn1 | 0.03 |
| Ty3_CRM | 0.69 |
| Ty3_Galadriel | 0.04 |
| Ty3_Tekay | 4.19 |
| Ty3_Reina | 0.80 |
| Non LTR elements |  |
| LINE | 0.96 |

|  |  |  |
| --- | --- | --- |
| ClassII |  |  |
|  | CACTA | 1.05 |
|  | hAT | 0.07 |
|  | MuDR_Mutator | 0.08 |
|  | Mariner | 0.01 |
|  | Helintron | 0.25 |
| Total |  | 42.83 |

The repetitive fraction corresponded to 42.83% of the *C. nitida* genome. It was possible to annotate 198 of the 330 clusters, the rest (6.56% of the total genome) remained unclassified (Table 2). The dispersed repetitive DNA sequences corresponded to 22.01% of the total genome. LTR-retrotransposons from the Ty1/Copia superfamily comprised 5.6% of the genome, while Ty3/Gypsy elements were 1.6 times more abundant (8.86%). Within Ty1/Copia, the SIRE lineage was the most abundant with 3.5%, while the Tekay lineage was the most represented among Ty3/Gypsy with 4.19%. LTR elements without a clear lineage classification corresponded to 5.13% of the total genome. LINEs corresponded to 0.96%, while Class 2 transposable elements corresponded to 1.46%, with CACTA being the most abundant (1.05%).

Among tandem repeats, the 5S rDNA showed a large abundance, comprising 4.75% of the *C. nitida* genome, even larger than the 35S rDNA (3.1%). Other tandem repeats (satDNA) corresponded to 6.4% of the genome. TAREAN identified six clusters with high-confidence satellites and six with low-confidence. Another six clusters, not identified by the TAREAN, showed typical circular graphs, and were confirmed as tandem repeats by DOTTER. The comparative dot-plot with the consensus sequences of the 18 identified satDNAs revealed some sequences with similarity to each other and were grouped into three superfamilies (Table 3; Fig. S2 and Fig. S3).

**Table 3:**
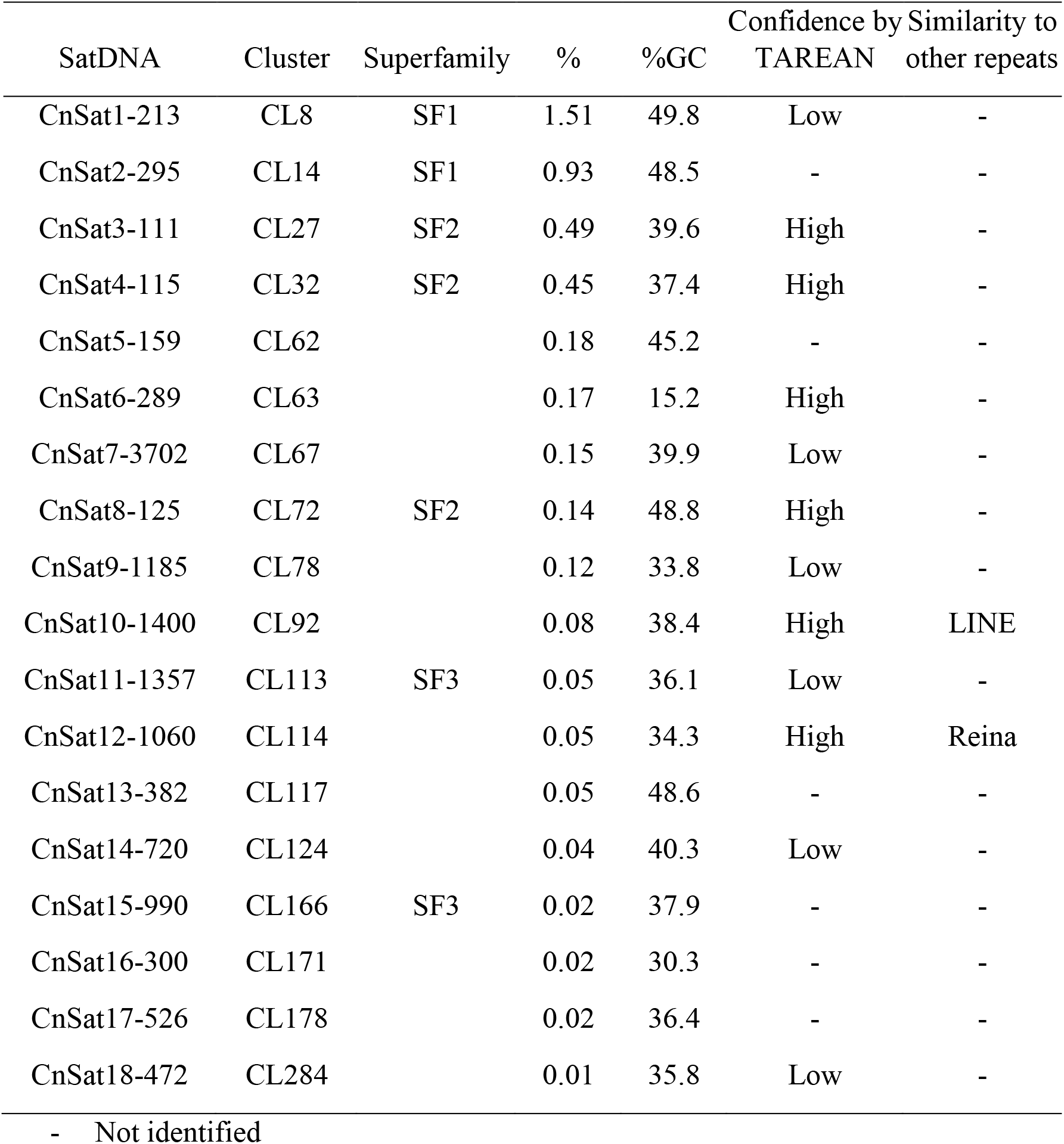
Satellite DNA families and superfamilies identified in *C. nitida* genome, showing genome proportion (%), percentage of Guanine and Citosine (%GC), TAREAN output and similarity to other repeats. First number in the name of the satellite represents its order of abundance, while the second number, the consensus size of its monomer sequence

Superfamily 1 (SF1), with 2.44% abundance, is composed of two satellites: CnSat1-213, classified by the TAREAN with 49.8% of GC, and CnSat2-295, not identified by the TAREAN, with 48.5% of GC showing 68.6% similarity between consensus sequences. The superfamily 2 (SF2) is composed of satellites classified with high-confidence by TAREAN, CnSat3-111, CnSat4-115 and CnSat8-125, showing 72.3% similarity among consensus sequences, lower average GC content (around 41.9%) and representing together 1.08% of the genome (Fig. 1). The third superfamily is composed of two satellites, one classified with low confidence and one not identified by TAREAN, CnSat11-1357 with 36.1% GC and CnSat15-990 with 37.9% GC. Together they corresponded to 0.07% of the genome (Table 3). The consensus sequences are provided in Supplementary Table 2. In addition to these satellites, some microsatellites were identified and described in Supplementary Table 3.

**Fig 1.**
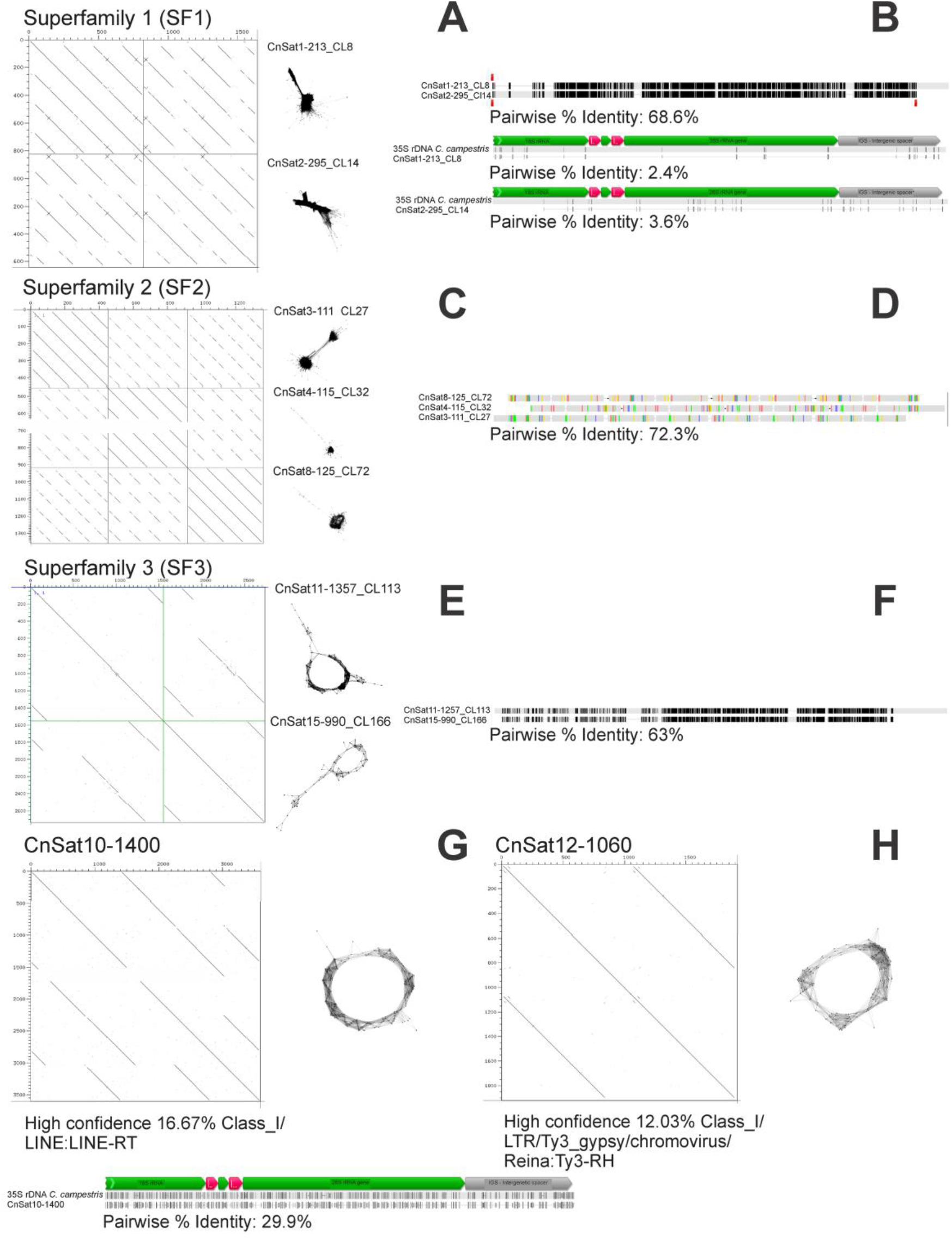
DOTTER and cluster graphics of hybridized satellites in *C. nitida*. **A** and **B,** Superfamily 1; **C** and **D,** Superfamily 2; **E** and **F,** superfamily 3; **G,** satellite CnSat10-1400; and **H,** CnSat12-1060. **B**, **D** and **F** show the alignments and similarity between the satellites subunits that constitute Superfamilies 1, 2 and 3, respectively. **B** and **G**, similarity of the subunits with the 35S rDNA previously assembled from *C. campestris* for Superfamily 1 and CnSat10-1400, respectively.

**Fig. 2.**
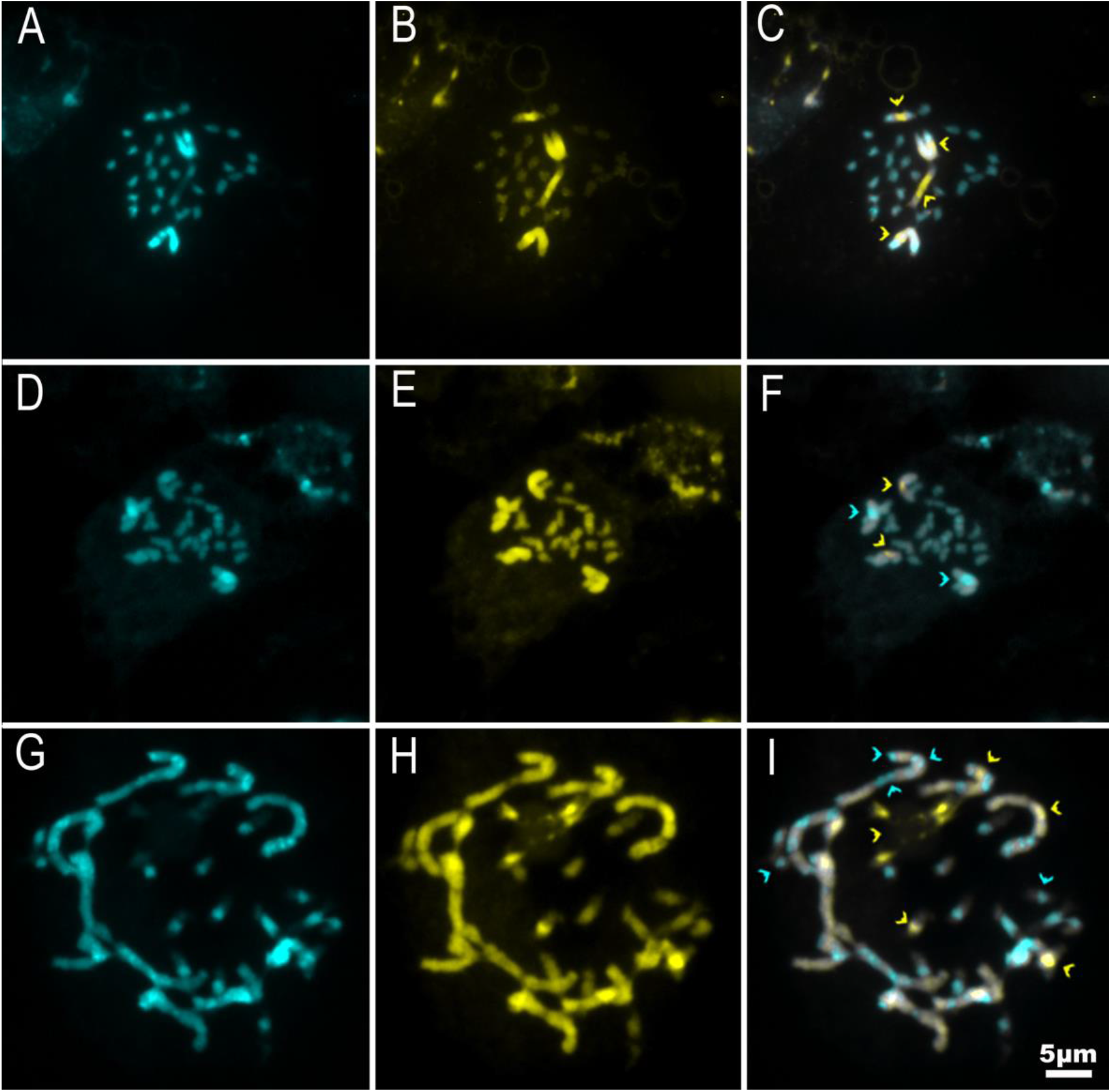
Metaphases of *C. nitida* (**A, B** and **C**), *C. africana* (**D, E** and **F**) and *C. angulata* (**G, H** and **I**) stained with CMA (yellow) and DAPI (blue). Overlapping in **C, F** and **I**. Arrowheads in **C, F** and **I** highlight heterochromatic bands in each karyotype.

**Fig. 3.**
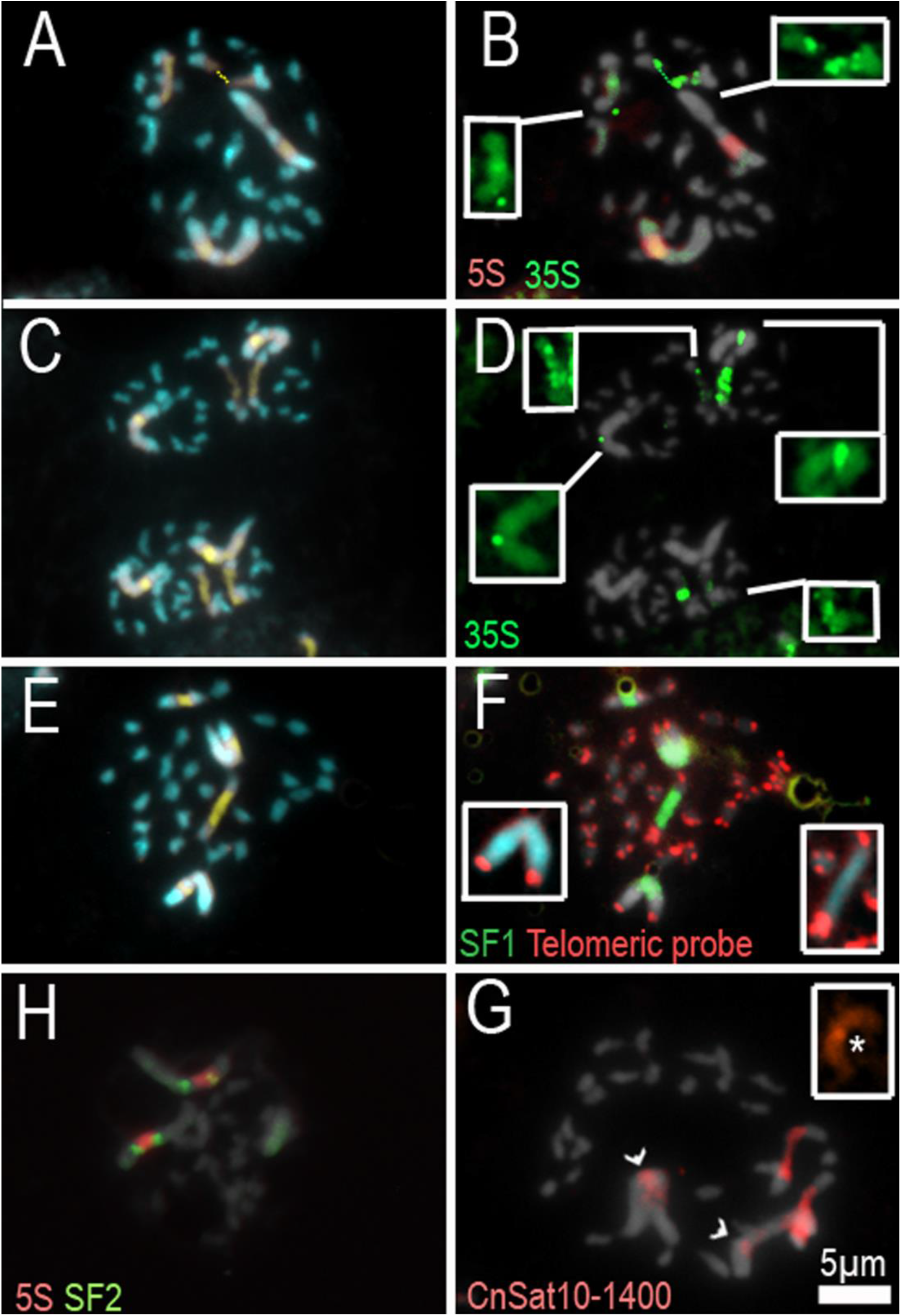
Metaphases of *C. nitida* showing colocalization of CMA^+^ bands (**A, C** and **E**) with 5S rDNA in red (**B**) and 35S rDNA in green (**B** and **D**) and SF1 superfamily in green (**F**). Insets in **B** and **D** show smaller 35S sites. In, **F**, the telomeric probe detected only terminal loci. In **H**, the 5S rDNA (in red) is flanked by the SF2 superfamily sites (in green). In **G**, CnSat10-1400 satellite signals in red; inset shows a detail of the satellite signal on one of the chromosomes of the largest pair. Chromosomes were counterstained with DAPI (blue, **A, C** and **E**, or grey).

Two satellites classified with high confidence showed low similarity to retrotransposons. CnSat10-1400 showed similarity with a LINE element (16.67% identity) and CnSat12-1060 showed similarity with a Ty1/Copia from the Reina lineage (12.03%). The consensus sequence of CnSat10-1400 was aligned against the reverse transcriptase (RT) domain of the LINE element, with 32.2% identity, while CnSat12-1060 showed 39.5% identity to the Ribonuclease H (RH) domain of Reina. Altogether, the repetitive fraction showed a more abundant and diverse tandem repeat fraction within *C. nitida* genome.

### Mapping repetitive sequences

Different repeats were selected for investigating their chromosomal distribution and putative association with heterochromatin and the largest chromosome pairs. Apart from the 5S and 35S rDNA, four others satDNA were selected: superfamily 1 (SF1), the most abundant among satDNAs, superfamily 2 (SF2), and two tandem repeats that showed similarity with mobile elements. Superfamily SF1 signals colocalized with 35S rDNA in both large chromosome pairs. The SF1 signals, however, were stronger and more extended than the rDNA signal, occupying the proximal region on the long arm of the largest pair (Fig. 3F). Superfamily SF2 is also located in the largest chromosome pair, presenting two signals on each homologue. These signals flanked the 5S rDNA sites (Fig. 3H). The 5S rDNA and SF2 sites occupy most of the short arm of the largest chromosome pair. The CnSat10-1400 satellite showed a small signal in the pericentromeric region of the short arm of the largest pair and a larger signal in the distended region of the second pair, similar to the 35S rDNA. (Fig. 3G). Despite colocalization of satellites SF1 and CnSat10-1400 with the 35S rDNA, these satellites did not show any significant *in silico* similarity with the 35S rDNA assembled from *C. campestris* (2.4% to CnSat1-213, 3.6% to CnSat2-295 and 29.9% to CnSat10-1400, identity to the aligned sequence, Fig. 1). CnSat12-1060, on the other hand, showed no evident chromosome hybridization (data not shown). The ND-FISH with telomeric probe showed terminal signals in all chromosomes of the complement, but no interstitial signals that could suggest previous chromosome fusions (Fig. 3F).

The most abundant LTR retrotransposon lineages, SIRE (Ty1/Copia), Tekay and Retand (Ty3/Gypsy), as well as the putative centromeric CRM lineage (Ty3/Gypsy), were also selected for hybridization in *C. nitida* chromosomes. The SIRE element showed signals along the entire length of the largest chromosome pair, with a gap in the 5S rDNA site. In addition, it also labelled the distal regions of the second largest pair, not including the proximal NOR (Fig. 4A). The Retand element showed a similar pattern, but it also displayed dispersed signals along the distended region of the NOR and along the 5S rDNA cluster (Fig. 4B). CRM labelled the largest chromosomes pairs, but also showed a weak labelling of the small chromosomes, slightly enriched in the pericentromeric region at least in some of them (Fig. 4C). Tekay element showed scattered proximal signals on the largest chromosome pairs and no signal in the NOR (Fig. 4D).

**Fig 4.**
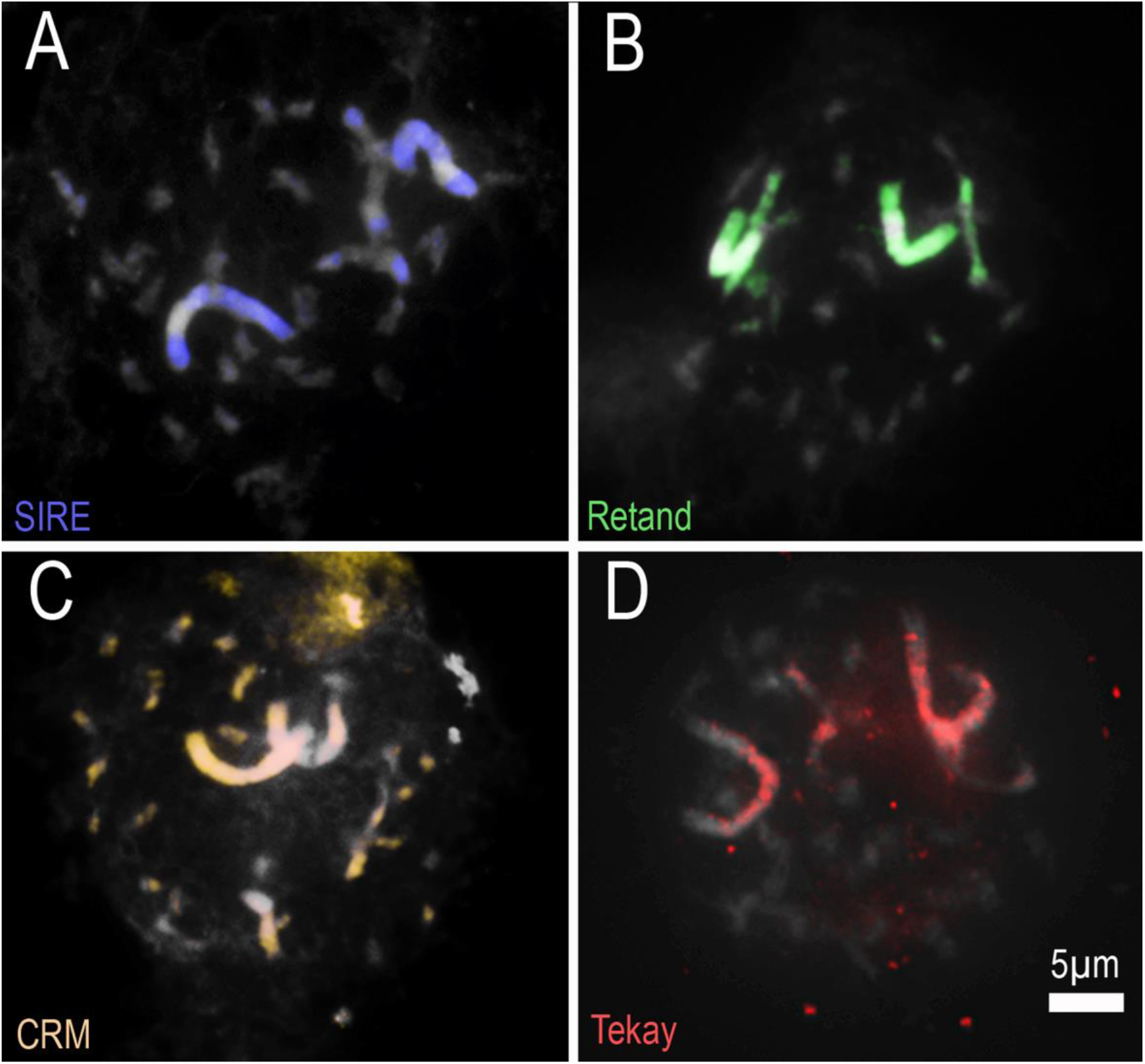
Metaphases of *C. nitida* showing the distribution of LTR Retrotransposons with an enrichment in the largest chromosome pairs. In **A**, the element Ty1/Copia SIRE (violet); in **B**, Ty1/Copia Retand (green); in **C**, Ty3/Gypsy CRM (yellow); and in **D**, Ty3/Gypsy Tekay (red). Chromosomes were counterstained with DAPI (grey).

Combined, these data demonstrate the enrichment of the large chromosome pairs of the *C. nitida* karyotype with tandem (rDNAs and satDNAs) and disperse (LTR retrotransposons) repetitive sequences. Several of these repeats colocalize in the largest pairs, evidencing a complex chromosome organization of this large chromosomes and indicating them as cause for the bimodal karyotype in this species.

## Discussion

All three analysed species - *C. nitida, C. africana*, and *C. angulata* - presented bimodal karyotypes. Although the remaining two species, *C. gerrardii* Baker and *C. natalensis* Baker, should be analysed in the future for confirmation, the presence of bimodal karyotype is likely a synapomorphy of *Cuscuta* subgenus *Pachystigma*. The phylogenetic relationships within the subgenus resolved *C. nitida* as sister to a clade with *C. natalensis* and *C. gerrardii* (García *et al*. 2014), with this clade sister to *C. africana* + *C. angulata*. This suggests that this karyotypic feature was maintained in the whole subclades, thus supporting the hypothesis that all species of this subgenus share bimodal karyotypes. In fact, the four subgenera of *Cuscuta* are not only delimited by phylogenetic, biogeographic, and morphological data, but each present unique cytogenetic peculiarities, such as the presence of holocentric chromosomes in the subgenus *Cuscuta* (García *et al*., 2014; Costea *et al*., 2015; García *et al*., 2019).

Much of the cytogenetic studies in the genus *Cuscuta* are restricted to conventional staining techniques with an emphasis on chromosome counting. Fewer studies conducted more detailed cytogenetic analyses, such as CMA/DAPI banding for characterization of heterochromatin. The latter studies revealed a numerical variation of these bands, with species having few bands, like *C. denticulata* Engelm. with only one pair of evident CMA^+^/DAPI^-^ bands, to species like *C. monogyna* Vahl, with approximately 90 CMA^+^/DAPI^-^ bands and 80 DAPI^+^/CMA^-^ bands in pachytene (Ibiapino *et al*., 2019, 2020). Despite this variation, our previous unpublished data showed that karyotypes with less bands are more frequent, even in polyploid species. In the majority of those cases, CMA^+^ bands are found in pericentromeric regions and are colocalized with 5S and 35S rDNA. In the species with a higher number of CMA^+^/DAPI^-^ or DAPI^+^/CMA^-^ bands, they are localised in interstitial regions. There is no evidence that this banding pattern is different for each subgenus and there may be similar patterns between different subgenera (Ibiapino *et al*., 2020). In the three species of the subgenus *Pachystigma* analysed here, the multiple CMA/DAPI bands were mainly present at the largest pairs. These bands do not differ much in number and position from those already reported in the genus, however, in these bimodal karyotypes, the bands are larger. In *C. nitida*, for example, the largest CMA^+^/DAPI^-^ band occupies a large part of the short arm of the largest chromosome pair. *Cuscuta africana* showed a similar pattern, however the largest heterochromatic band was DAPI^+^/CMA^-^. The large number of heterochromatic bands on the smallest chromosomes is not observed in the other two species of the *Pachystigma* subgenus. This characteristic may indicate an incipient accumulation of heterochromatin in these chromosomes, which could eventually lead to a less asymmetrical karyotype, such as the amplification observed in other unrelated *Cuscuta* species, such as *C. indecora* Choisy (*Grammica* subgenus) and *C. monogyna* (*Monogynella* subgenus) (Ibiapino *et al*., 2019), indicating that this character is homoplastic in the genus. Alternatively, karyotype asymmetry in *Pachystigma* may be maintained by an unknown mechanism.

Bimodality in *Pachystigma* is not due to chromosome fusion. Although *C. nitida* has 2*n* = 28, lower than the basic number proposed for the genus *Cuscuta*, which is *x* = 15 (Pazy and Plitmann, 1995), the sizes of the two largest pairs cannot be explained by a single fusion of two pairs of small chromosomes. In addition, the ND-FISH with telomeric probe did not provide evidence for any interstitial sites in *C. nitida*, which may indicate that there was no fusion event in the origin of this karyotype. Furthermore, *Cuscuta angulata* presented 2*n* = 30, showing no reduction in chromosome number and a bimodal karyotype. Many *Cuscuta* species have 2*n* = 30, but there are species with 2*n* = 8, 10, 14, 16, 18, 20, 28, 30, 32, 34 and polyploids with 2*n* = 28, 42, 44, 56, 60, 90, 150. *Cuscuta epithymum* (L.) L. (subgenus *Cuscuta*), for instance, shows an intraspecific variation which could be attributed to chromosome fusions and polyploidy, with 2*n* = 14, 16, 28, 30, 32 and 34. Individuals with 2*n* = 14 and 2*n* = 32 are bimodal, while 2*n* = 16 and 2*n* = 34 are symmetric (García and Castroviejo, 2003; García *et al*., 2019). Three species of subgenus *Cuscuta* (holocentric), *C. epithymum* (2*n* = 14), *C. europaea* L. (2*n* = 14), and *C. epilinum* Weihe (2*n* = 6*x* = 42), had a reduction in the chromosome number. In the case of this subgenus, there may have been chromosomal fusion events, since holocentric chromosomes have diffuse kinetochores, and consequently these chromosomes can stabilize fragments or fused chromosomes favouring rearrangements (Mandrioli and Manicardi, 2020).

Bimodal karyotypes may also originate through interspecific hybridization, as proposed for the genus *Agave*, in which allopolyploid species might have chromosomes of different sizes inherited from different parents (McKain *et al*., 2012). In *Cuscuta*, there are numerous cases of interspecific hybridization and polyploidy (reviewed by García *et al*., 2014). For example, *C. veatchii* Brandegee is an allopolyploid originated from the hybridization of *C. denticulata* and *C. nevadensis* I.M. Johnst. With 2*n* = 60, *C. veatchii* possess 30 smaller chromosomes and 30 slightly larger chromosomes with very evident centromeres, characteristic of *C. denticulata*, and *C. nevadensis*, respectively (Ibiapino *et al*., 2019). However, molecular phylogenetic analyses have showed that reticulate evolution occurs mainly in the subgenus *Grammica* (e.g., Stefanović and Costea, 2008; Costea and Stefanović, 2010; García *et al*., 2014; Costea *et al*., 2015a). There is also preliminary phylogenetic evidence suggesting that some species of subgenus *Cuscuta* may have a hybrid origin. Different accessions of *C. approximata*, for example, have polymorphism in the ITS, and the location of *C. kurdica* differed between the ITS and *trnL* trees (García and Martín, 2007). These contrasting topologies may indicate hybridization events similar to those reported in subgenus *Grammica* (e.g., Stefanović and Costea, 2008; García *et al*., 2014). Evidence such as this has not been observed in *C. africana, C angulata*, and *C. nitida* with ITS, 26S, *trnL* nor *rbcL* sequences analyses (García and Martín, 2007; García *et al*., 2014). Chromosome number and size, as well as the number of rDNA sites in *C. nitida*, were within the range of variation already reported for species of the genus *Cuscuta*. So far, most *Cuscuta* species have shown few rDNA sites, varying from two to 36 sites of 5S rDNA and from two to 30 sites of 35S rDNA (Fogelberg, 1938; García, 2001; García and Castroviejo, 2003; Guerra and García, 2004; McNeal *et al*., 2007; Ibiapino *et al*., 2019, 2020; García *et al*., 2019). Thus, our results do not suggest neither hybridization nor polyploidy as the cause of bimodality in *Pachystigma*.

*Cuscuta africana* presented divergence in chromosome number compared to previous report (2*n* = 30, García *et al*., 2019). Intraspecific variation is unlikely because samples were plants collected from the same population. It is more likely that the 2*n* = 30 reported earlier was a mistake, since conventional staining may leave the proximal, distended NOR unnoticed. NORs are more easily identified as CMA^+^/DAPI^-^ bands (see, for example Fig. 2D and 2F). Similar miscounts have been registered for *Passiflora foetida* L., which was first described as having 2*n* = 22 (Snow and MacDougal, 1993) and later corrected to 2*n* = 20 (De Melo and Guerra, 2003).

Because chromosome fusions and intraspecific hybridization seem less probable, repetitive sequences accumulation in specific chromosome pairs could be a probable mechanism for karyotype asymmetry in subg. *Pachystigma*. Indeed, repetitive DNA in *Cuscuta* is involved in the expansion of the genome, causing an increase in chromosomes, such as in *C. monogyna* and *C. indecora* (Ibiapino *et al*., 2020; Neumann *et al*., 2021). In these two cases, however, chromosomes increased proportionally in size, maintaining karyotype symmetry, and resulting in similar karyotypes, although *C. monogyna* and *C. indecora* belong to different subgenera (*Monogynella* and *Grammica*, respectively). All 12 *Cuscuta* species sequenced by Neumann *et al*. (2021) showed a greater abundance of LTR type elements, with SIRE being the most dominant among Ty1/Copia lineages and Tekay most dominant among Ty3/Gypsy, ranged from 8.5% to 30.8% for Ty1/Copia and 7.3% to 28.5% for Ty3/Gypsy. Many small genome species such as *C. pentagona* Engelm. showed a higher proportion of Class II elements, representing 12.6% of the genome. A large fraction of these genomes was also composed of satellite DNA, reaching up to 18% in *C. europaea*. In this species, the satDNA CUS-TR24 is the major constituent of its heterochromatic bands (Vondrak *et al*., 2021). Similar results were observed for *C. nitida*, with 3.5% SIRE and 4.19% Tekay. Satellites also made up a significant percentage of the *C. nitida* genome, 6.4%. However, Class II elements showed a low proportion, 1.46%. These demonstrate that repetitive accumulation is a common mechanism in the evolution of genomes within *Cuscuta* genus, increasing the size of chromosomes within a particular karyotype.

The evident accumulation of repetitive DNA sequences in the largest chromosomal pairs of *C. nitida* supports the influence of heterochromatin in the karyotype asymmetry of *Cuscuta*. In the bimodal karyotypes of subgenus *Pachystigma*, the most evident heterochromatic bands are restricted to the largest pairs. The 5S rDNA corresponded to 4.75% of the genome of *C. nitida* and was colocalized with the largest CMA^+^ band of that species. In addition, all hybridized satellite DNAs, as well as the 35S rDNA sites and most of the transposable elements, are restricted or highly enriched in the largest chromosomal pairs (Fig. 5). In *Muscari* Mill. (Asparagaceae), a massive amplification of the MCSAT satDNA family occurred in only one chromosome pair. This single satDNA family corresponds to 5% of the total genome of *M. comosum* (L.) Mill. and contributed to the progressive increase in the karyotype asymmetry of *Muscari* species (de la Herrán *et al*., 2001). In *Eleutherine*, two of the *E. bulbosa* satellites, Ebusat1 and Ebusat4, occur in the interstitial region of the largest pair of *E*. *bulbosa* and *E. latifolia*, both with bimodal karyotypes. In addition, the four most abundant retrotransposons also showed accumulation in the larger pair. This demonstrates that accumulation of repetitive sequences can generate an increase of only part of the chromosomes of a karyotype and lead to a change in karyotype symmetry (Báez *et al*., 2019). This suggests that the bimodality of *Pachystigma* subgenus could also originate from the asymmetric expansion of multiple repetitive DNA lineages (Fig. 5).

**Fig. 5.**
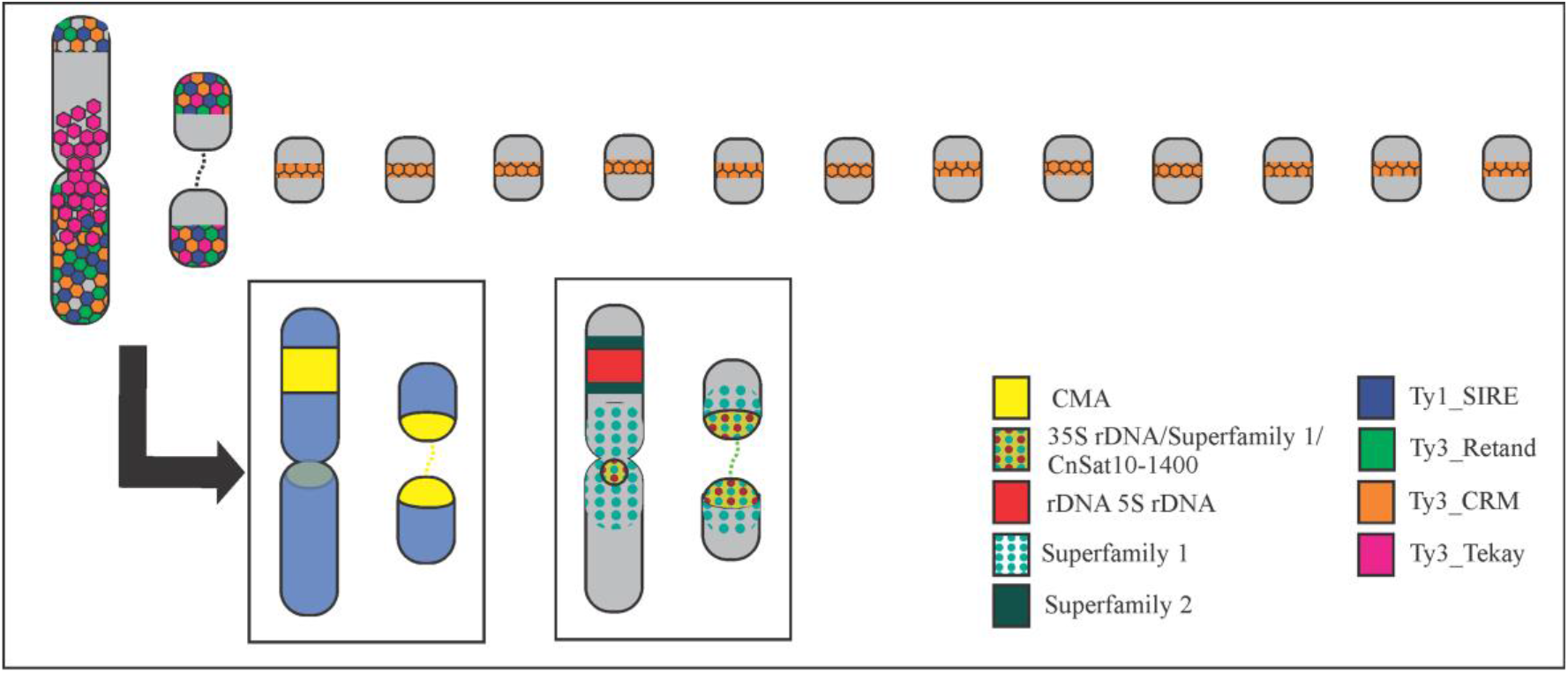
Schematic representation of the distribution of heterochromatic bands and repetitive sequences in *C. nitida* bimodal karyotype. Ribosomal DNAs (5S e and 35S) and satellite superfamily SF1 colocalized with CMA^+^ heterochromatin in the largest chromosome pairs, with SF1 and SF2 also adjacent to it. LTR-retrotransposons were enriched in the largest pairs, except for the CRM element, also present in the smallest chromosomes.

The SF1 and CnSat10-1400 signals colocalized with the 35S rDNA but showed no similarity with the 35S rDNA of *C. campestris*, which is a species of subg. *Grammica*. This may suggest that this satellite DNA unit has originated from tandem duplications of a less-conserved, intergenic region of *Cuscuta* rDNA, such as the IGS, or that it was inserted in *C. nitida* rDNA locus after the divergence between subgenera *Grammica* and *Pachystigma*. In *Phaseolus* L. (Fabaceae), jumper satDNA was inserted into the NTS region of 5S rDNA (Ribeiro *et al*., 2017*b*). However, the 35S rDNA and CnSat10-1400 formed independent clusters, suggesting that this satDNA has not become part of the rDNA unit but is rather interspersed along the rDNA site. Nevertheless, no 35S rDNA cluster from *C. nitida* presented a circular graph, indicating its completeness, marking it impossible to confirm at present if it is part of the rDNA unit of this specific species. Furthermore, CnSat10-1400 showed higher similarity with the reverse transcriptase domain of a LINE element, which may indicate a possible origin of this satellite from a TE, and later interspersion within the 35S rDNA loci, or the insertion of LINEs in this satDNA, as observed in *Cuscuta europaea* for CUS-TR24 (Vondrak *et al*., 2021).

The satellites of *C. nitida* CnSat10-1400 and CnSat12-1060 showed similarity with transposable elements, LINE and Reina, respectively. Some transposable elements and repetitive genes can contribute to the formation and dissemination of satellite DNAs. In *Lathyrus sativus* L., most of the satellites originated from small tandem repetitions present in the 3’ untranslated region of the Ogre retrotransposons (Vondrak *et al*., 2020). MITE transposable elements were appointed as generators of satellite DNA in bivalve molluscs and *Drosophila*. Similarly, in ants of the genus *Messor*, a Mariner element gave rise to the expansion of satellite DNA IRE-130 (Palomeque and Lorite, 2008). In fishes, copies of the 5S rDNA originated the satellite 5SHindIII. Ancestors of tRNA were probably responsible for the formation of tandemly repeated sequences in higher plants (López-Flores and Garrido-Ramos, 2012). In humans, it has been identified that a quarter of all mini/satellites are derived from transposable elements. TE-derived satellites usually have monomers above the standard 500 bp of size and generally occupy pericentromeric regions (Meštrović *et al*., 2015). This is the case of CnSat10-1400 and CnSat12-1060, with monomers of 1,400 bp and 1,060 bp, respectively. In *Pisum sativum*, variants of the satellite PisTR-A are incorporated into Ty3/Gypsy Ogre elements. The untranslated region that separates the 3’ gag-pol domains from the LTR is highly variable in the pea Ogre elements and carries several other tandem repeats (Macas *et al*., 2009). In maize, the CRM1TR and CRM4TR tandem repeats are entirely derived from centromeric retrotransopsons (CM) (Sharma *et al*., 2013). None of the satellites found in *C. nitida* showed similarity to satellites previously described by Oliveira *et al*. (2020) in *C. europaea*, a species of the subgenus *Cuscuta*, indicating that these satellites have independent origins and the composition of heterochromatic bands in holocentric and monocentric chromosomes of the genus or between different subgenera are different.

## Supplementary data

**Table S1**. Primer pairs designed for probe amplification for FISH.

**Table S2**. Consensus sequences of the satellites described in *C. nitida*.

**Table S3**. Motif and proportion of microsatellites in *C. nitida* genome.

**Fig. S1.** Histogram showing the distribution of analysed reads in *C. nitida* clusters and superclusters.

**Fig. S2.** Dotplots and cluster graphics of all remaining satellites annotated in *C. nitida* genome. Some satellites were not detected by TAREAN, but the dot plot and cluster graphics characterised them as putative tandem repeats.

**Fig. S3.** Comparative dot plot of all satellites described *C. nitida*. This dot plot was used for identifying similarities between different satellites and define satellite superfamilies.

## Conclusions

The well-supported clade of subgenus *Pachystigma* is characterized by the presence of bimodal karyotypes in all species analysed. Although the three species had different CMA/DAPI band patterns, these bands were more enriched in the larger chromosomes of the three karyotypes. The genome organization of *C. nitida* repetitive fraction suggested a differential chromosome accumulation of diverse repetitive families, mainly satDNA, rDNA and retrotransposons, as the probable mechanism of origin for the bimodal karyotypes within this subgenus. This shows that the increase in chromosomes, which led to the emergence of bimodality in this clade, was associated to the accumulation of repetitive sequences in heterochromatin. The composition of this heterochromatin may be different among species. In *Cuscuta*, the amount and diversity of repetitive DNA is high and satellite DNAs can originate from transposable elements and potentially be incorporated or interspersed with the rDNA.

## Acknowledgments

We thank the Fundação de Amparo a Ciencia e Tecnologia de Pernambuco (FACEPE) for the financing of the Postgraduate scholarship; the Conselho Nacional de Desenvolvimento Científico e Tecnológico (CNPq) and the Coordenação de Aperfeiçoamento de Pessoal de Nível Superior (CAPES, Financial Code 001) for the financial support for the development of the project. Computational resources for the Galaxy based RepeatExplorer were provided by the ELIXIR-CZ project (LM2015047), part of the international ELIXIR infrastructure. The field work was supported in part by the Natural Sciences and Engineering Research Council of Canada Discovery grant (no. 326439).

## Author contribution

AI, MAG, and AHP designed the research. SS provided materials. AI and MB performed the *in silico* analysis. AI performed the banding characterization and *in situ* hybridizations. AI and AHP performed the data interpretation and wrote the first draft of the manuscript. All co-authors participated in manuscript writing, revisions, and editing.

## Notes

### Competing Interest Statement

The authors have declared no competing interest.

